# Chronic *Staphylococcus aureus* lung infection correlates with proteogenomic and metabolic adaptations leading to an increased intracellular persistence

**DOI:** 10.1101/414409

**Authors:** Xin Tan, Mathieu Coureuil, Elodie Ramond, Daniel Euphrasie, Marion Dupuis, Fabiola Tros, Julie Meyer, Ivan Nemanzny, Cerina Chhuon, Ida Chiara Guerrera, Agnes Ferroni, Isabelle Sermet-Gaudelus, Xavier Nassif, Alain Charbit, Anne Jamet

**Affiliations:** Université Paris Descartes, INSERM U1151 - CNRS UMR 8253, Institut Necker-Enfants Malades. Team: Pathogenesis of Systemic Infections, Paris, France; Plateforme Métabolomique Institut Necker-Enfants Malades, Structure Fédérative de Recherche SFR Necker, University Paris Descartes, Paris, France; Plateforme Protéome Institut Necker-Enfants Malades, PPN, Structure Fédérative de Recherche SFR Necker, University Paris Descartes, Paris, France; Proteomics platform 3P5-Necker, Université Paris Descartes - Structure Fédérative de Recherche Necker, INSERM US24/CNRS UMS3633, Paris, France; Laboratoire de Microbiologie de l’hopital Necker, University Paris Descartes, Paris, France; Université Paris Descartes, INSERM U1151 - CNRS UMR 8253, Institut Necker-Enfants Malades. Team: Canalopathies épithéliales: la mucoviscidose et autres maladies, Paris, France

**Author notes:** **Corresponding author:** Anne Jamet. **Alternate corresponding author:** Alain Charbit; Bâtiment Leriche. 14 Rue Maria Helena Vieira Da Silva CS 61431 - 75993 PARIS – FRANCE, —.

**Keywords:** Cystic fibrosis, *Staphylococcus aureus*, intracellular persistence, biofilm, proteogenomics

## Abstract

**Background:** Chronic lung infection of cystic fibrosis (CF) patients by *Staphylococcus aureus* is a well-established epidemiological fact. Indeed, *S. aureus* is the most commonly identified pathogen in the lungs of CF patients. Strikingly the molecular mechanisms underlying *S. aureus* persistency are not understood.

**Methods:** We selected pairs of sequential *S. aureus* isolates from 3 patients with CF and from one patient with non-CF chronic lung disease. We used a combination of genomic, proteomic and metabolomic approaches with functional assays for in-depth characterization of *S. aureus* long-term persistence.

**Results:** For the first time, we show that late *S. aureus* isolates from CF patients have an increased ability for intracellular survival in CFBE-F508del cells compared to ancestral early isolates. Importantly, the increased ability to persist intracellularly was confirmed for *S. aureus* isolates within the own patient F508del epithelial cells. An increased ability to form biofilm was also demonstrated.

Furthermore, we identified the underlying genetic modifications inducing altered protein expression profiles and notable metabolic changes. These modifications affect several metabolic pathways and virulence regulators that could constitute therapeutic targets.

**Conclusions:** Our results strongly suggest that the intracellular environment might constitute an important niche of persistence and relapse necessitating adapted antibiotic treatments.

**Summary:** *S. aureus* persists for years in the lungs of patients with cystic fibrosis despite antibiotic therapies. We demonstrate that *S. aureus* adaptation leads to increased intracellular persistence suggesting a key role for intracellular niche during *S. aureus* chronic lung infection.

## Introduction

*Staphylococcus aureus* and *Pseudomonas aeruginosa* are the most common pathogens infecting the lungs of patients with a chronic lung disease including cystic fibrosis (CF) [1, 2]. Furthermore, *S. aureus* is one of the earliest bacteria detected in infants with CF. However, very few studies have addressed the adaptations undergone by *S. aureus* in this context [3, 4].

*S. aureus* has the ability to form biofilm [5-7] and to survive within a wide range of eukaryotic host cells [8-17]. These abilities are likely to contribute to the persistence of *S. aureus* in airways of patients with chronic lung diseases despite appropriate antimicrobial treatments [18, 19]. *S. aureus* persistence is associated with a drastic decrease in metabolism [20], a decrease in the expression of virulence factors and an increase in the expression of bacterial adhesins [21]. Such profile is typical of small-colony variants (SCVs) that are defined by small-sized colonies [15, 22, 23]. Beside SCVs, strains with normal colony morphology can exhibit similar patterns of “low toxicity” which allow them to persist intracellularly without being cleared by host cell defense mechanisms [21]. A “low toxicity” pattern can be achieved either transiently, following changes in the expression of genes encoding toxins and/or regulators, or permanently, by mutations in global regulators [24-26].

By studying serial isolates, we show that, during long-term lung infection, *S. aureus* adaptation occurs through genomic modifications that accumulate over time and lead to major metabolic modifications and protein expression changes. We also reveal that persistence of *S. aureus* is associated with convergent evolution responsible for an increased ability to form biofilm as well as to survive within host cells. These observations should be taken into account in therapeutic decisions aiming at eradicating *S. aureus* chronic infections by choosing drugs specifically targeting biofilm-embedded and intracellular bacteria.

## Methods

Whole genome sequencing was performed on an Illumina MiSeq instrument (2×150 bp) and the sequences were processed using the Nullarbor bioinformatic pipeline software v1.20 and RAST server. The sequences reported in this paper are available at NCBI’s BioProject database under accession number PRJNA446073 (http://www.ncbi.nlm.nih.gov/bioproject/446073).

Quantification of biofilm formation was assessed with crystal violet staining in polystyrene 96-well plates. Cystic Fibrosis Bronchial Epithelial cell line CFBE41o- and primary nasal epithelial cell were infected with a multiplicity of infection of 100 using an inoculum taken from cultures of *S. aureus* grown in Brain Heart Infusion until exponential growth phase. Infected cells were kept for 6 days in a medium containing 50 μg/mL gentamycin to kill extracellular bacteria. For proteomics, proteins were digested and analyzed by liquid chromatography coupled with tandem mass spectrometry (nanoLC-MS/MS). For metabolomics, metabolite profiling of *S. aureus* isolates was performed by liquid chromatography–mass spectrometry (LC-MS). The mass spectrometry proteomics data have been deposited to the ProteomeXchange Consortium via the PRIDE [27] partner repository with the dataset identifier PXD011281.

A full description of methods is available in Supplementary Methods.

### Statistical analysis

Data were analyzed using R or GraphPad Prism softwares. Results are presented either with one representative experiment for clarity or as means ± standard deviation (SD). The number of biological and technical replicates is indicated per figure.

For two-sample comparisons statistical significance was measured using unpaired two-tail Student’s t-test or Wilcoxon rank sum test as indicated in the figure legends. For comparisons between more than two groups, statistical significance was measured using one-way analysis of variance (ANOVA) with multiple comparisons (Dunnett’s correction) performed, with each value compared to that of the reference strain.

P values of <0.05 were considered to indicate statistical significance.

### Ethics statement

All experiments were performed in accordance with the guidelines and regulations described by the Declaration of Helsinki and the low Huriet-Serusclat on human research ethics and informed consent was obtained for all participating subjects. Serial isolates of *S. aureus* were obtained from airway secretions from four patients with chronic lung infection at the Necker-Enfants Malades University Hospital, Paris, France. Sputum sampling is part of routine standard care. The research procedure is validated by Ile de France 2 IRB (ID-RCB/Eudract: 2016 A00309-42).

## Results

### Selection of *S. aureus* sequential isolates from patients with chronic lung infection

Three patients with CF (CF1, CF2 and CF3) and, for comparison purpose, one patient with non-CF chronic lung disease (CLD) were chosen. For each patient we selected one early and one late isolate separated by 3 to 9 years intervals. Whole-genome sequencing confirmed that each pair of isolates belonged to four distinct clones (**Figure 1**). Patient diseases and treatments are detailed in supplementary Methods.

**Fig 1.**
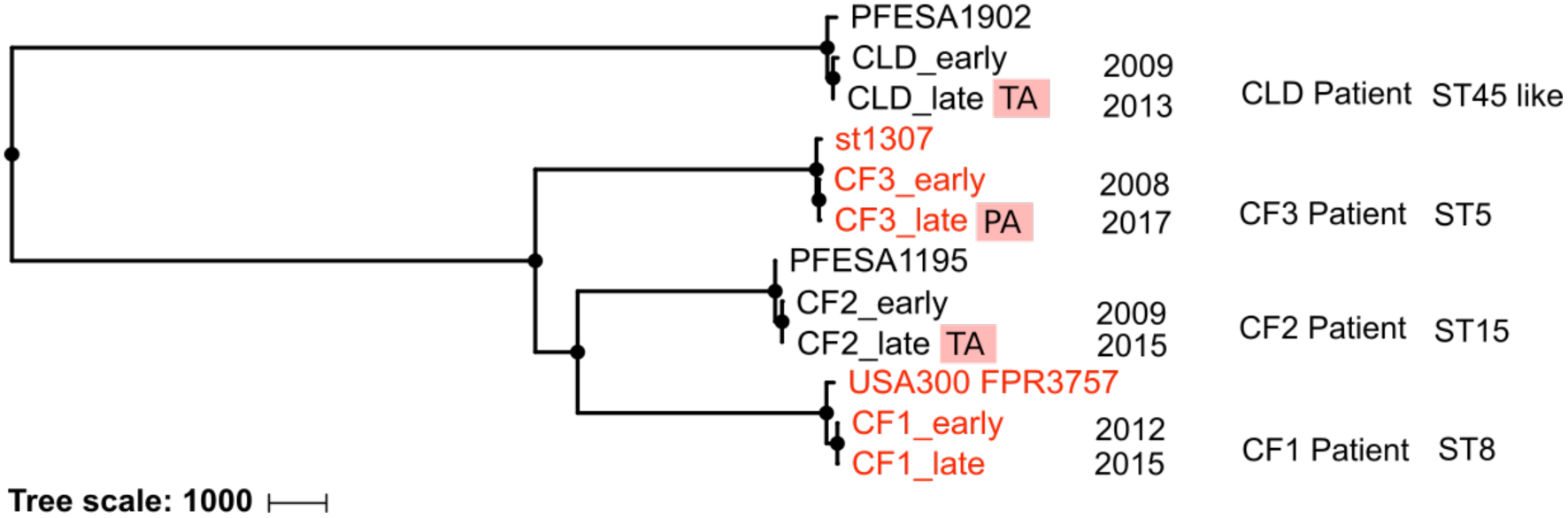
Selection of four pairs of *S. aureus* isolates belonging to four different STs in four patients. Dendrogram generated by wgsa.net from the genomes of the eight clinical isolates retrieved from the respiratory samples of three patients with CF, one patient with non-CF chronic lung infection and four reference genomes from public databases. Branch length is proportional to the number of variant nucleotide sites within the core genes. For each patient, the isolate taken first is called “early” while the isolate taken later is named “late”. The dates of sampling and the sequence type (ST) of the isolates are indicated. “TA” and “PA” mean that the isolate is auxotrophic for thymidine or pantothenate respectively. The name of the isolate is indicated in red when it is resistant to methicillin (MRSA). Reference strains included are PFESA1902 (ERR554197), st1307 (ERR158691), PFESA1195 (ERR554722).

### *S. aureus* clinical isolates from CF patients evolved an increased persistence ability within CFBE-F508del epithelial cell line

Numerous studies have shown that *S. aureus* has the ability to survive within human cells [8-17]. We subsequently aimed at investigating if during the course of within-lung adaptation, *S. aureus* isolates have evolved a greater ability to persist within epithelial cells. We infected bronchial CFBE epithelial cell line (F508del +/+ CFTR mutation) with clinical isolates, the control strain USA300-LAC and a stable SCV mutant altered in the haemin biosynthetic pathway (hereafter named Δ*hem*). As expected, wild-type bacteria were not able to persist whereas the Δ*hem* mutant was able to persist intracellularly during the whole course of the experiment (**Figure S1**) [13]. All early and late clinical isolates were able to persist at least 2.6-fold, and up to 900-fold, more than the USA300-LAC reference strain at day 3 and 6 post-infection (**Figure 2AB**). Furthermore, at day 3 and 6 post-infection, all the late isolates recovered from CF patients exhibited an improved ability to persist intracellularly within CFBE-F508del epithelial cells compared to cognate early isolates (**Figure 2A**). Interestingly, the CLD_late isolate recovered from the non-CF patient did not exhibit an improved ability to persist within CFBE epithelial cells compared to CLD_early isolate (**Figure 2B**). These data suggest that *S. aureus* adaptation within CF-lungs correlates with an improved ability to persist intracellularly in cells with a CFTR dysfunction.

**Fig 2.**
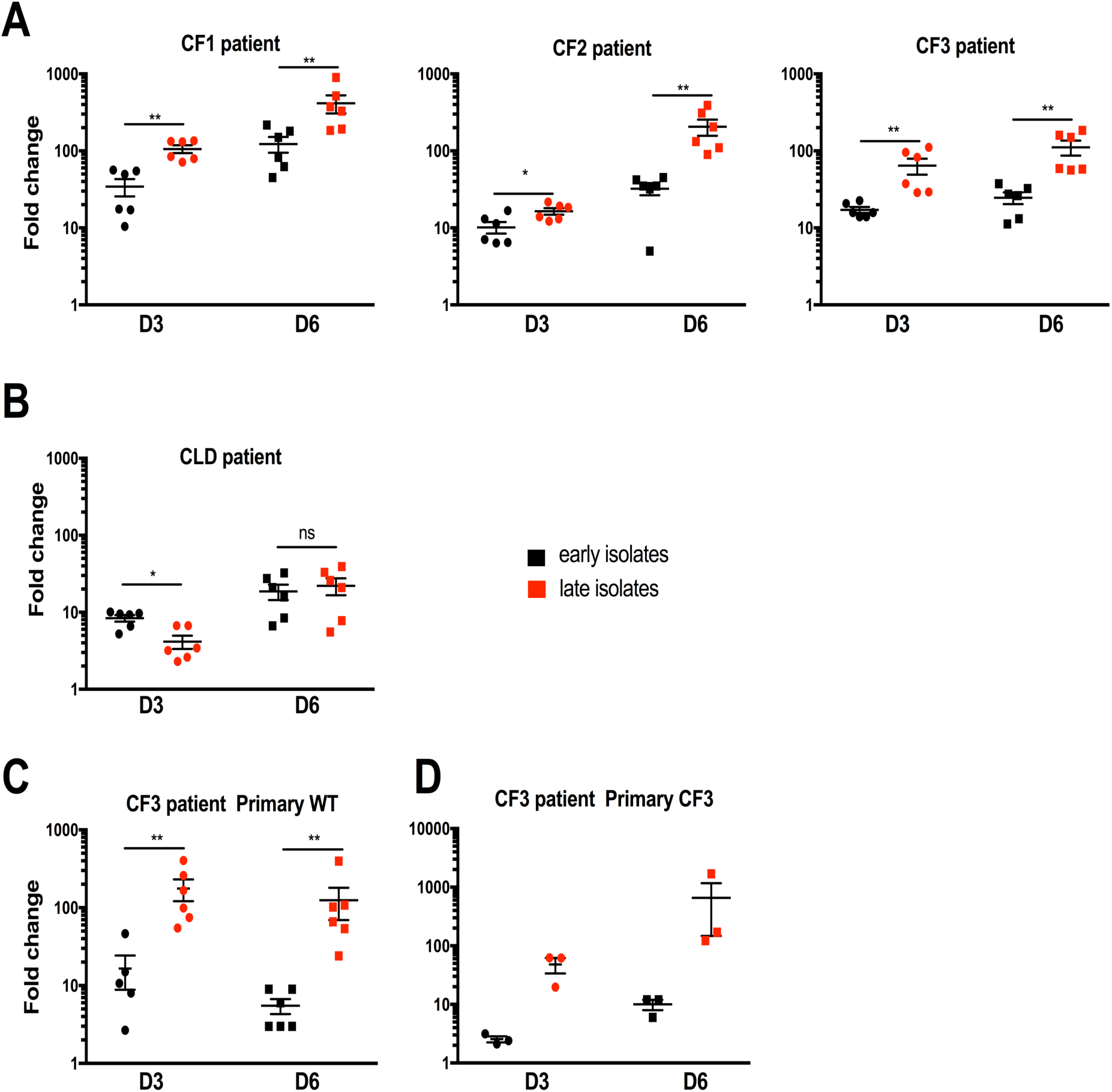
Intracellular persistence of *S. aureus* clinical isolates in CFBE-F508del epithelial cell line and within primary patient cells. **A and B)** Bronchial CFBE epithelial cell line (CFBE-F508del homozygous for the F508del-CFTR mutation) was infected with the control strain USA300-LAC and clinical isolates from CF patients (A) or CLD patient (B). **C and D)** Primary nasal epithelial cells retrieved from a healthy donor (“Primary WT”) (C) and from the CF3 patient (“Primary CF3” with F508del +/+ CFTR mutation) (D) were infected with the control strain USA300-LAC and CF3 isolates. For all experiments, gentamicin was present throughout the experiment to prevent extracellular bacterial growth and new infection. Bacterial loads inside cells were evaluated by CFU enumeration at 3 and 6 days after infection. Results are normalized with USA300- LAC strain as a reference and expressed as a fold change of CFUs. Results have been obtained from two independent experiments performed in triplicate for ABC and one experiment for D. Statistical analysis was performed by Wilcoxon rank sum test *P < 0.05; **P<0.01; ns P>0.05.

### Late *S. aureus* isolate of CF3 patient exhibits an increased persistence within primary F508del epithelial own patient cells

To confirm the relevance of the results obtained with bronchial CFBE epithelial cell line, we first assessed the persistency of CF3 isolates within primary epithelial cells isolated from the nose of a healthy donor (**Figure 2C**). In addition, we performed an infection assay with the CF3 primary epithelial own patient cells (F508del +/+ CFTR mutation) to verify the specific within patient-adaptation of *S. aureus* recovered from long-term infection (**Figure 2D**). These experiments confirmed that the late isolate persistence ability is improved compared to early isolate at day 3 and 6 within both primary nasal epithelial cells retrieved from a healthy donor and from the CF3 patient.

### *S. aureus* clinical isolates from chronically infected patients evolved high biofilm formation ability

Assuming that isolates retrieved from chronic infections might have a high biofilm-forming capacity, we studied the biofilm formation ability of the pairs of isolates. Remarkably, all isolates displayed a greater capacity to form biofilms compared to that of the weak biofilm-producer USA300-LAC reference strain (p-value of <0.001, **Figure S2**). Furthermore, for three chronically infected patients, the late isolates formed more biofilm than the early isolates, revealing that long-term adaptation within lungs had improved their biofilm formation capacity (p-value of <0.001, **Figure 3**).

**Fig 3.**
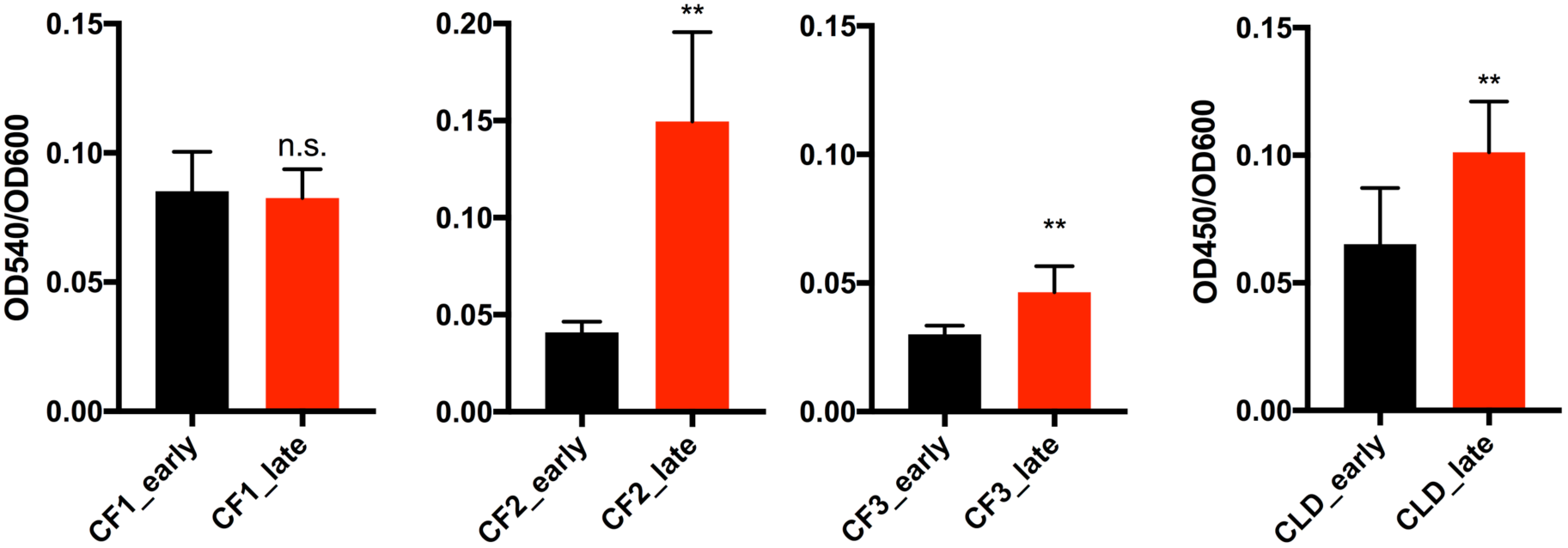
Quantification of biofilm formation of *S. aureus* clinical isolates. Biofilm formation quantification was performed using the crystal violet microtiter assay in BHI medium with 1% glucose. Results shown are the mean ±SD for three independent experiments performed in triplicate. Statistical significance was measured using a two-tail Student’s t-test when biofilm production of a late isolate was compared with biofilm production of cognate early isolate from the same patient. ** indicates p-value of <0.001 whereas ns indicates p-values>0.05.

### Late *S. aureus* clinical isolates from chronically infected patients acquired auxotrophies

Compared to that of USA300-LAC, all patients early isolates and the CF1 late isolate display similar colony morphology on brain heart infusion (BHI) agar plates and a wild-type growth in a liquid broth mimicking sputum (Cystic Fibrosis Sputum Medium or CFSM) **(Figure S3 and Figure 4A)**. In contrast, the late isolate of CF2 and CLD patients displayed a typical SCV phenotype with very small colonies on BHI agar (**Figure S4**) and CF2_late, CF3_late and CLD_late isolates exhibited a growth defect in CFSM broth **(Figure 4BCD)**. Thymidine-dependent SCVs are frequently isolated from patients treated with sulfamethoxazol (SXT) [28]. Indeed, supplementation with thymidine restored almost wild-type growth for CF2_late and CLD_late isolates (**Figure 4BD**). We used genomic data to determine the auxotrophy of CF3_late isolate and identified a frameshift in *panB* gene, which is involved in de novo biosynthesis of pantothenic acid (**Table 1**). Accordingly, growth of CF3_late isolate in the presence of pantothenate restored wild-type growth (**Figure 4C**). Thus, isolates from three out of four patients with *S. aureus* chronic lung infection acquired auxotrophy during the course of the disease.

**Fig 4.**
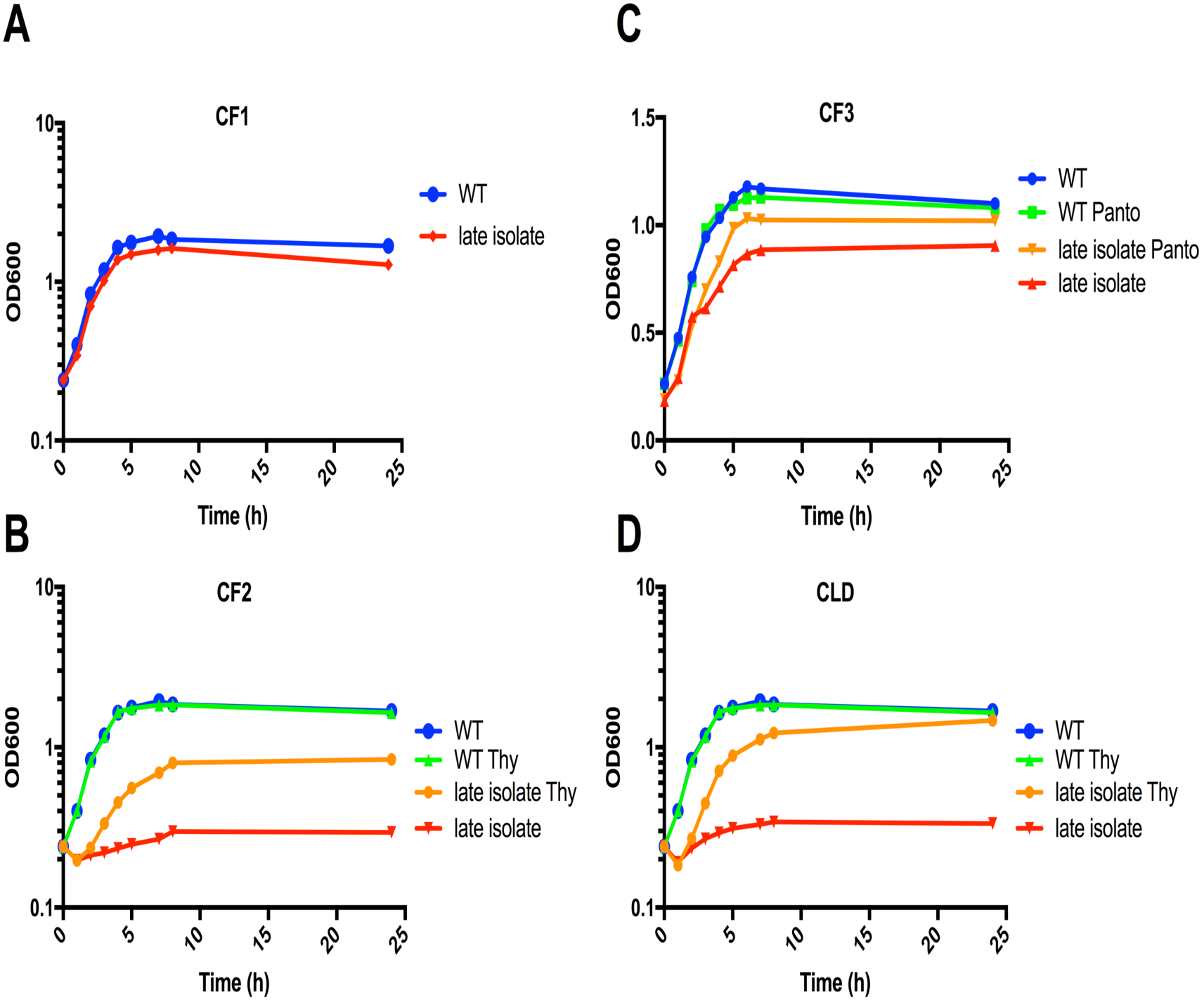
Growth of late *S. aureus* clinical isolates in CFSM. Growth curves were carried out in medium mimicking the respiratory fluid of cystic fibrosis patients (Cystic Fibrosis Sputum Medium or CFSM), with or without the addition of thymidine or pantothenate. The results shown correspond to a representative experiment. The orange and green curves correspond to bacterial growth in media supplemented with either thymidine or pantothenate; the red and blue curves, to bacterial growth in medium without thymidine or pantothenate. WT, USA300- LAC.

**Table 1.**
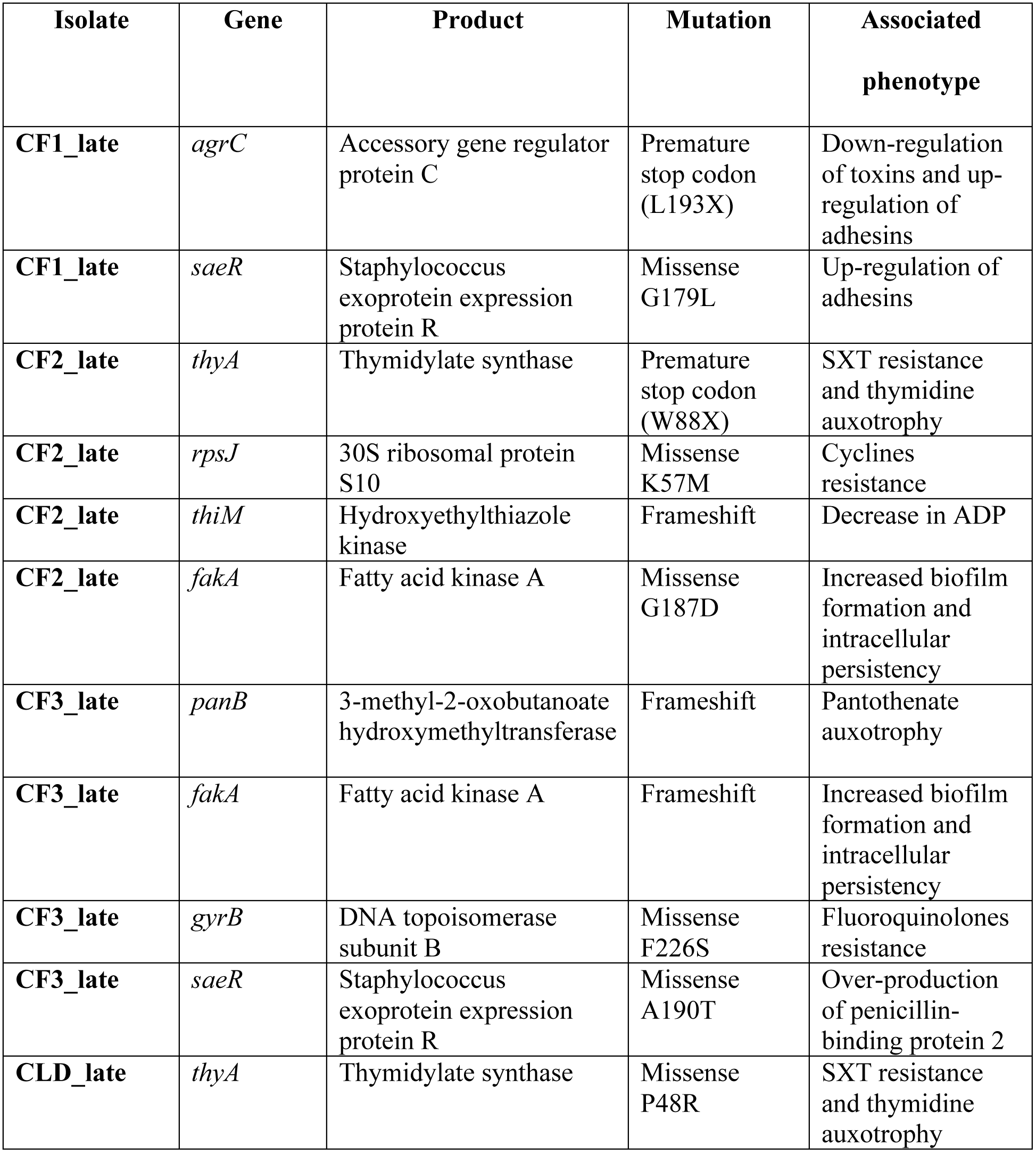
Mutations linked to phenotypic changes in clinical isolates.

### Late *S. aureus* clinical isolates from chronically infected patients acquired antibiotic resistance

Genome analysis evidenced mutations in *thyA*, *gyrB*, and *rpsJ* genes, which were associated with SXT, fluoroquinolones and cyclines resistance, respectively (**Table 1**). Thus, isolates from three out of four patients with *S. aureus* chronic lung infection acquired antibiotic resistance consistent with the administration of the corresponding drugs during the course of the disease.

### Genomic, proteomic and metabolomic modifications associated with *S. aureus* adaptation during chronic lung infection

In order to investigate the underlying genomic, proteomic and metabolic modifications associated with the observed phenotypic changes we compared genomes, proteomes and metabolomes of late compared to early isolates. The differences in proteomic and metabolic profiles between early and late isolates of patients are highlighted by heatmaps shown in **Figure S5**.

Genomes of all clinical isolates were de novo assembled and coding DNA sequences (CDSs) were annotated. Most of the SNPs were missense variants occurring in CDSs (**Table 2)**. Nonsynonymous mutations acquired by late isolates were found mainly in genes involved in metabolic processes (**Figure 5A**) and more specifically in “amino acid transport and metabolism” and “carbohydrate transport and metabolism” functional categories. In addition, the largest category of proteins to be differentially expressed for all pairs also comprised proteins related to metabolism processes (and more specifically to the “amino acid transport and metabolism” category) (**Figure 5B**). Concordant with genomic and proteomic results, the category “amino acids” was the most altered metabolites category in the late isolate of all patients compared to their cognate early isolates (**Figure 5C**).

**Fig 5.**
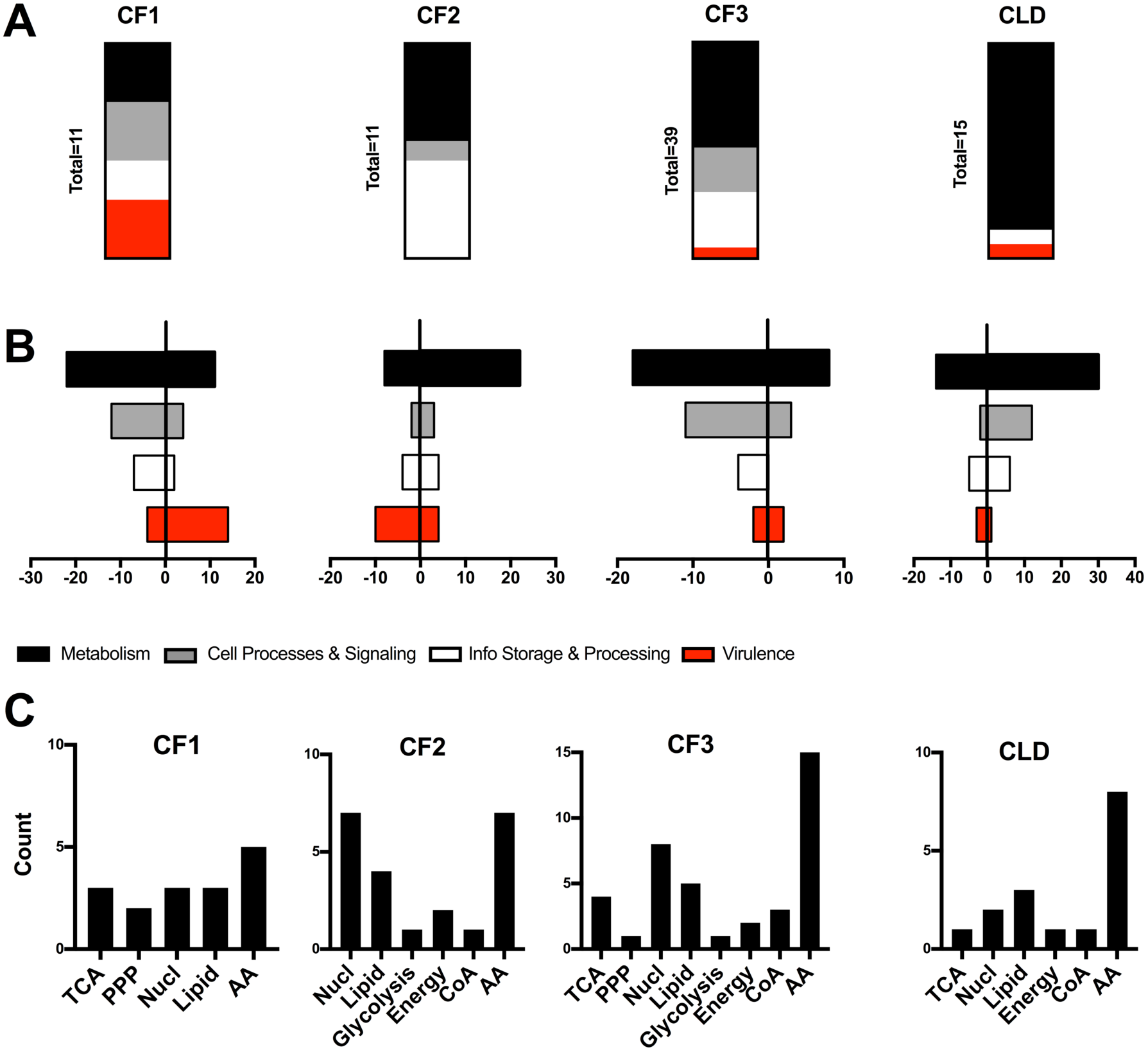
Proteogenomic and metabolomic analysis of the four pairs of *S. aureus* isolates. **A)** Vertical histograms show the functional classification of proteins encoded by genes with nonsynonymous mutations in the genomes of late isolates of *S. aureus* compared to early isolates. **B)** Horizontal histograms show the functional classification of differentially expressed annotated proteins in late compared to early isolates of each patient. For each category, histograms represent the number of down- and up-regulated proteins from proteomic analysis using the threshold of <2 and >2, respectively. Only genes and proteins with functional annotation available are included. The “cellular processes and signaling” category encompasses regulatory proteins and proteins involved in cell wall and capsule synthesis. The “information storage and processing” category encompasses proteins involved in replication, translation and repair processes. The “metabolism” category encompasses proteins involved in metabolism and transport. The “virulence” category encompasses exotoxins, proteins involved in adhesion, biofilm formation and immunomodulation. **C)** Categorization in 8 categories of altered amount of metabolites in late compared to early isolates. Metabolites were detected by carrying out 2 independent experiments performed in triplicate. TCA, Tricarboxylic acid cycle; PPP, Pentose phosphate pathway; Nucl, Nucleotides; CoA, Coenzyme A; AA, Amino acids.

**Table 2.** General features of detected mutations.

|  | **CF1_late** | **CF2_late** | **CF3_late** | **CLD_late** |
| --- | --- | --- | --- | --- |
| **Time since early isolate** | 2.8 years | 6.7 years | 9 years | 4.4 years |
| **Total polymorphisms** | 21 | 34 | 79 | 30 |
| **SNP<sup>a</sup>** | 19 | 25 | 73 | 26 |
| **INDEL<sup>b</sup>** | 2 | 9 | 6 | 4 |
| **CDS<sup>c</sup>** | 18 | 21 | 62 | 23 |
| **NON-SYN<sup>d</sup> (%)** | 14 (77.8) | 16 (47.1) | 51 (64.6) | 17 (56.7) |
| **FR<sup>e</sup>** | 2 | 4 | 4 | 2 |
| **MS<sup>f</sup>** | 10 | 11 | 47 | 14 |
| **STOP gained<sup>g</sup>** | 1 | 1 | 0 | 1 |
| **Other** | 1 | 0 | 0 | 0 |
| **SYN<sup>h</sup>** | 4 | 5 | 11 | 6 |
| **IG<sup>i</sup>** | 3 | 13 | 17 | 7 |

<sup>a</sup>SNP, single nucleotide polymorphism; <sup>b</sup>INDEL, insertion-deletion; <sup>c</sup>CDS, coding sequence;
<sup>d</sup>NON-SYN, nonsynonymous mutation; <sup>e</sup>FR, frameshift variant; <sup>f</sup>MS, missense variant;
<sup>g</sup>STOP gained, premature stop codon; <sup>h</sup>SYN, synonymous mutation; <sup>i</sup>IG, intergenic

Many regulatory proteins were differentially expressed. Indeed, proteins of the Agr, Rot, Sae, Sar or Fur regulatory networks were differently expressed in all late isolates. In CF1_late isolate, the *agrC* and the *saeR* genes mutations **(Table 1)** had a pleiotropic effect on the proteome (down-regulation of delta hemolysin and PSMb1 and upregulation of proteins encoded by *spa, sbi, fnbA, rot* and *coa* genes). In addition, adhesins encoded by *sasG, efb, sdrD* and *ecb* were up-regulated. In CF2_late isolate, the *agr* regulon is also downregulated suggesting an evolution toward low virulent and highly adhesive properties. The metabolite profiling of CF2_late isolate, showed a decrease in ADP, which is well correlated with the lack of ThiM (hydroxyethylthiazole kinase) expression found in proteomic analysis due to a frameshift in *thiM* gene **(Table 1)**. In CF3_late isolate, frameshifts in *fakA* and *panB* genes **(Table 1)** were associated with a lack of cognate proteins expression in CF3_late isolate. In addition, adhesins encoded by *sdrD* and *sasF* were upregulated. Interestingly, an over-production of penicillin-binding protein 2 encoded by *mecA* is correlated with the *saeR* mutation **(Table 1)** [29]. The metabolite profiling of CF3_late isolate, revealed a drastic diminution of pantothenate, coenzyme A and dephospho-coenzyme A, which is in line with the lack of expression of PanB and PanC proteins [30].

Of note, the non-CF control clone displays a different evolution trajectory. Indeed, AgrA and AgrC were up-regulated in CLD_late isolate, suggesting that it has retained virulent properties.

Altogether, the proteogenomic data suggest that all the late isolates recovered from CF patients (but not CLD patient) have evolved toward highly adhesive and low virulent properties. Besides, metabolic profiling suggests that all late isolates have evolved a reduced citric acid cycle activity compared to cognate early isolates.

## Discussion

Our study showed that during chronic lung infection, *S. aureus* adapts through the acquisition of common adaptive traits including antibiotic resistances, auxotrophies, reduced citric acid cycle activity, increased biofilm and intracellular persistence abilities that occurred irrespective of the clone type.

Of particular interest, we report mutations in two master regulatory systems, Agr and Sae, likely to impact multiple proteins expression and metabolites amounts.

*agr*-defective mutants, such as CF1 late isolate, have been shown to arise during chronic infections and are better adapted to persistence within the infected host [25, 31, 32].

Genetic alterations directly or indirectly targeting SaeR regulon were identified in the 3 CF patients. Since SaeR is involved in the regulation of over 20 virulence factor genes [33] and SaeRS-deficient bacteria are less infective in animal models [34], it is likely that SaeRS is a key factor in long-term colonization. In the three CF late isolates, we observed an increased in the expression of the SdrD adhesin belonging to the SaeR regulon and involved in adhesion to human nasal epithelial cells and to human keratinocytes [35][36]. Our results suggest that SdrD is also important for long-term lung colonization.

In patients with chronic lung infections, SCVs detection is most often the consequence of a long-term SXT treatment [37]. Mutations in the *thyA* gene, as found in CF2 and CLD late isolates, lead to stable clinical SCVs that are no longer susceptible to SXT and are thymidine-auxotrophic (TA-SCV) [28, 37]. Since thymidine is assumed to be abundant during lung inflammation, TA-SCVs can still grow in this environment.

In CF3 late isolate, we observed a pantothenate auxotrophy, which has been previously associated with persistency in *Mycobacterium tuberculosis* [38]. The acquisition of pantothenate auxotrophy suggests that pantothenate could also be present in CF lungs. Thus, our data confirm that metabolic specialization is a common phenomenon among long-term colonizers [39].

Other striking traits of phenotypic convergent evolution of *S. aureus* identified in this work were the increased ability to form biofilm and to persist in the intracellular niche. For CF2 and CF3 patients, the increased biofilm ability of late isolates could be linked to a mutation in the *fakA* gene, encoding fatty acid kinase A (FakA). Indeed, several studies showed that FakA-null strains were proficient in biofilm formation [6] and deficient in the expression of virulence factors controlled by the SaeRS system [40]. Overexpression of adhesins detected in proteomic analysis could also ultimately lead to increase biofilm formation in clinical isolates.

Numerous studies have demonstrated *S. aureus* ability to persist within host cells [8-17]. Strikingly, for the three CF patients, the *S. aureus* late isolates showed a greater ability to persist within CFBE-F508del epithelial cells compared to the early ones at day 3 and 6 post-infection. Of note, the late isolate of CLD patient did not present an improved ability to persist intracellularly within CFBE-F508del epithelial cells possibly due the fact that it has adapted to a non-CF patient.

Our multi-omics approach allowed both confirmation of previously known mechanisms and identification of novel candidate genes and pathways involved in the persistence ability of clinical isolates. We now provide evidence that the *saeR*/*fakA* regulon and the pantothenate pathway could also be promising therapeutic targets to fight persistent *S. aureus* infections.

Our study suggests that the use of antibiotic with a good intracellular penetration should be the best therapeutic option in order to eradicate *S. aureus* from chronically infected lungs.

## Funding

This work was supported by Institut national de la Santé et de la Recherche Médicale; Centre National de la Recherche Scientifique; and Université Paris Descartes Paris Cité Sorbonne. A scholarship from the China Scholarship Council [n° CSC NO. 201508500097] has been provided to XT.

The funders had no role in study design, data collection and analysis, decision to publish, or preparation of the manuscript.

## Acknowledgments

We thank Aurélie Hatton, Charlotte Roy and Zhicheng Zhou for their help with primary nasal epithelial cells culture.

## Author contributions

XT, MC, XN, AC and AJ conceived and designed the study. XT, ER, MD, DE, FT, JM and AJ made the experiments and analysis. IN, CC, ICG, AF and ISG contributed with data and analysis. AJ, AC and XT wrote the manuscript, with contributions and comments from all authors.

**No conflicts exist for the authors**

## Supplementary figure

**Figure S1.**
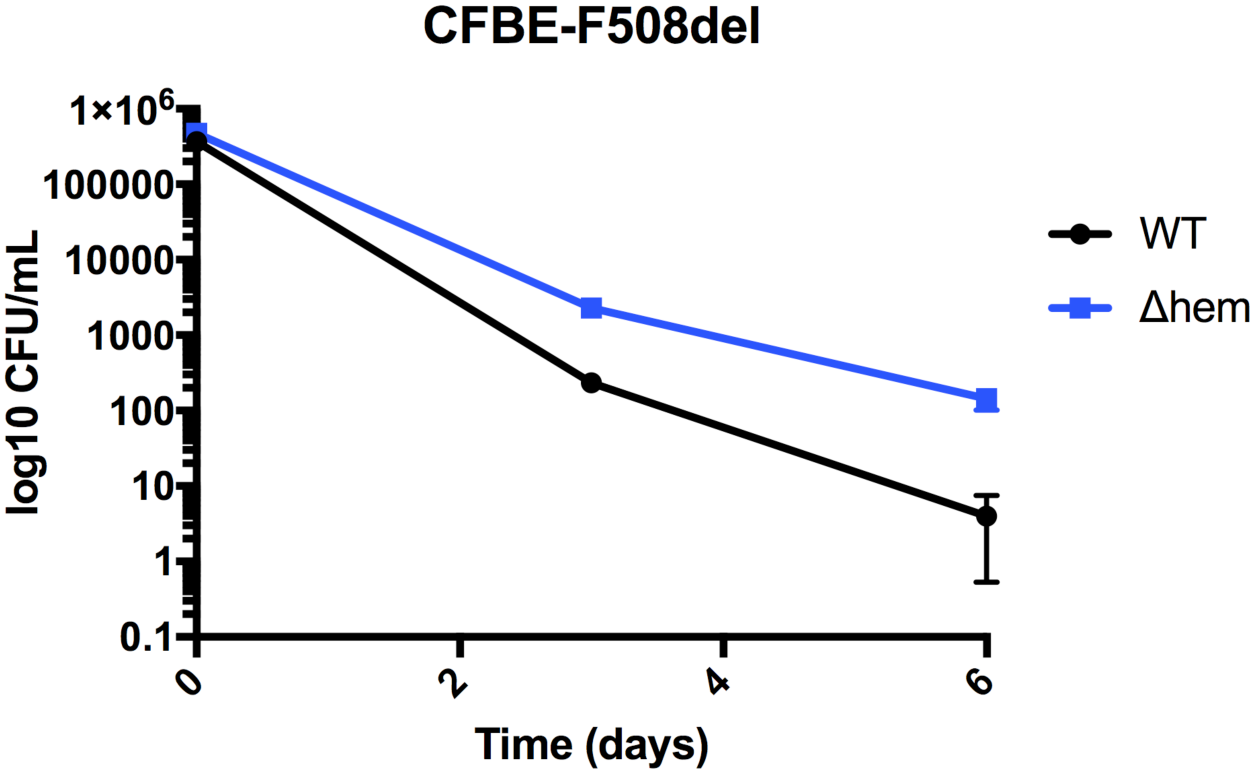
Intracellular growth curves of reference *S. aureus* isolates in CFBE epithelial cell line. Bronchial CFBE epithelial cell line (CFBE-F508del homozygous for the F508del-CFTR mutation) was infected with the control strain USA300-LAC (WT) and a stable SCV mutant altered in the haemin pathway (Δ*hem*). Gentamicin was present throughout the experiment to prevent extracellular bacterial growth and new infection. Bacterial load inside cells were evaluated by CFU enumeration at 3 and 6 days after infection. Results shown are the mean ±SD for four experiments performed in triplicate.

**Figure S2.**
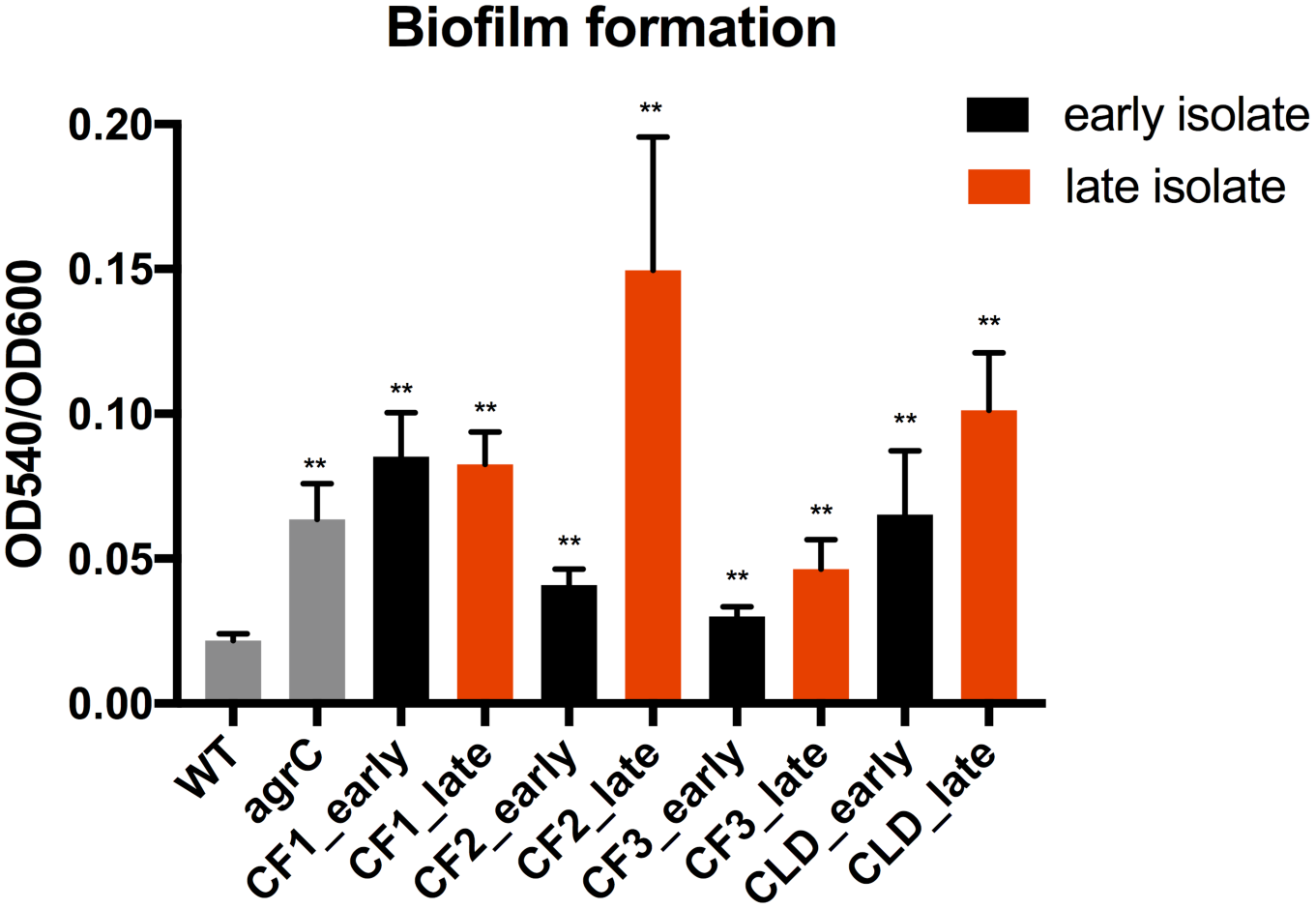
Quantification of biofilm formation of *S. aureus* clinical isolates. Biofilm formation was monitored using the crystal violet microtiter assay in BHI medium with 1% glucose. USA300-LAC strain (WT) was used as a reference for weak biofilm production whereas its *agrC* derivative obtained from the Nebraska Transposon Mutant Library was used as a reference for strong biofilm production. Results shown are the mean ±SD for three independent experiments performed in triplicate. Statistical significance was measured using one-way ANOVA with multiple comparisons (Dunnett’s correction) performed on the dataset as a whole, with each value compared to the WT. ** indicates p-value of <0.001.

**Figure S3.**
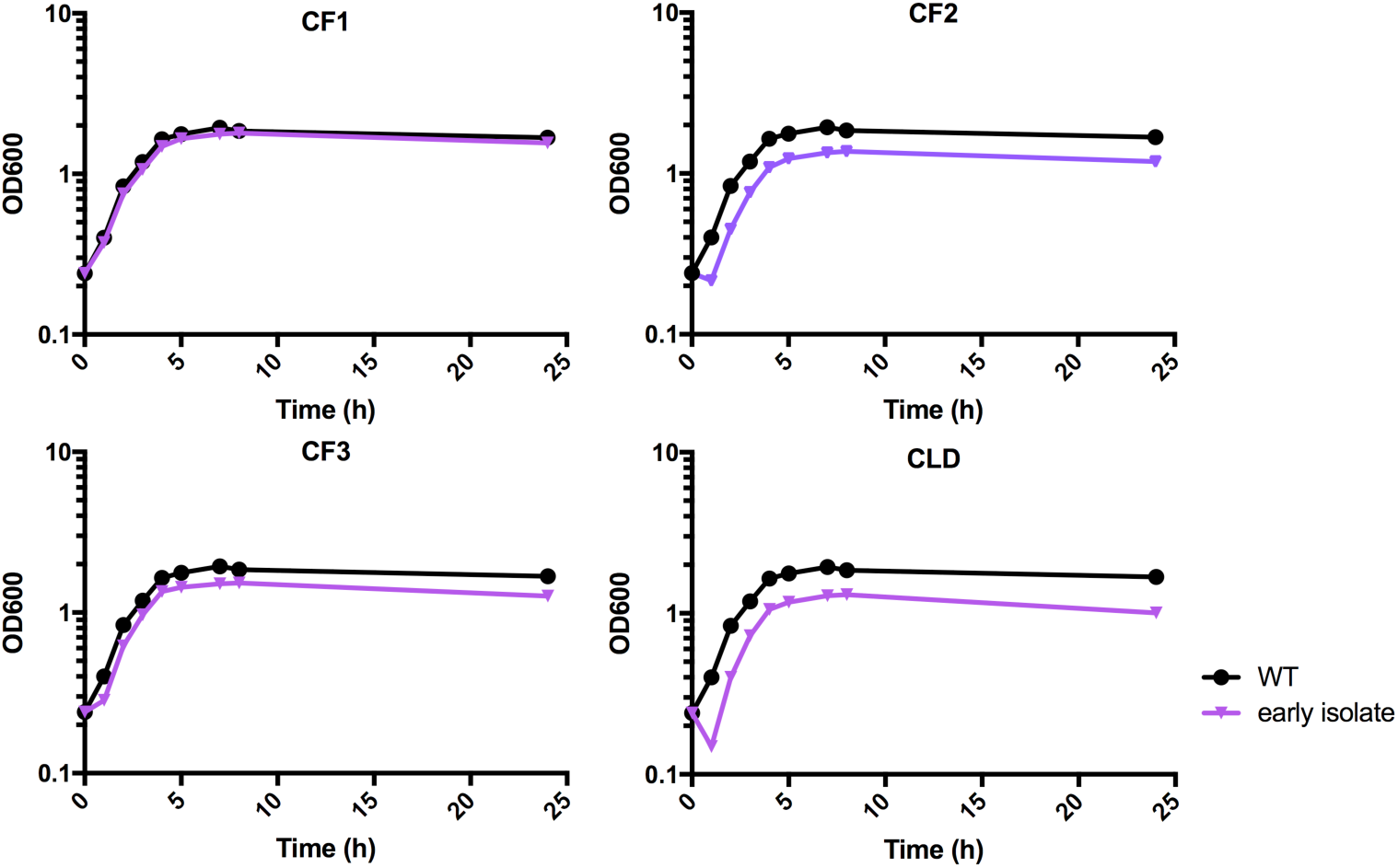
Growth curves of early *S. aureus* clinical isolates in CFSM. Growth curves were carried out in CFSM. The results shown correspond to a representative experiment. WT, USA300-LAC.

**Figure S4.**
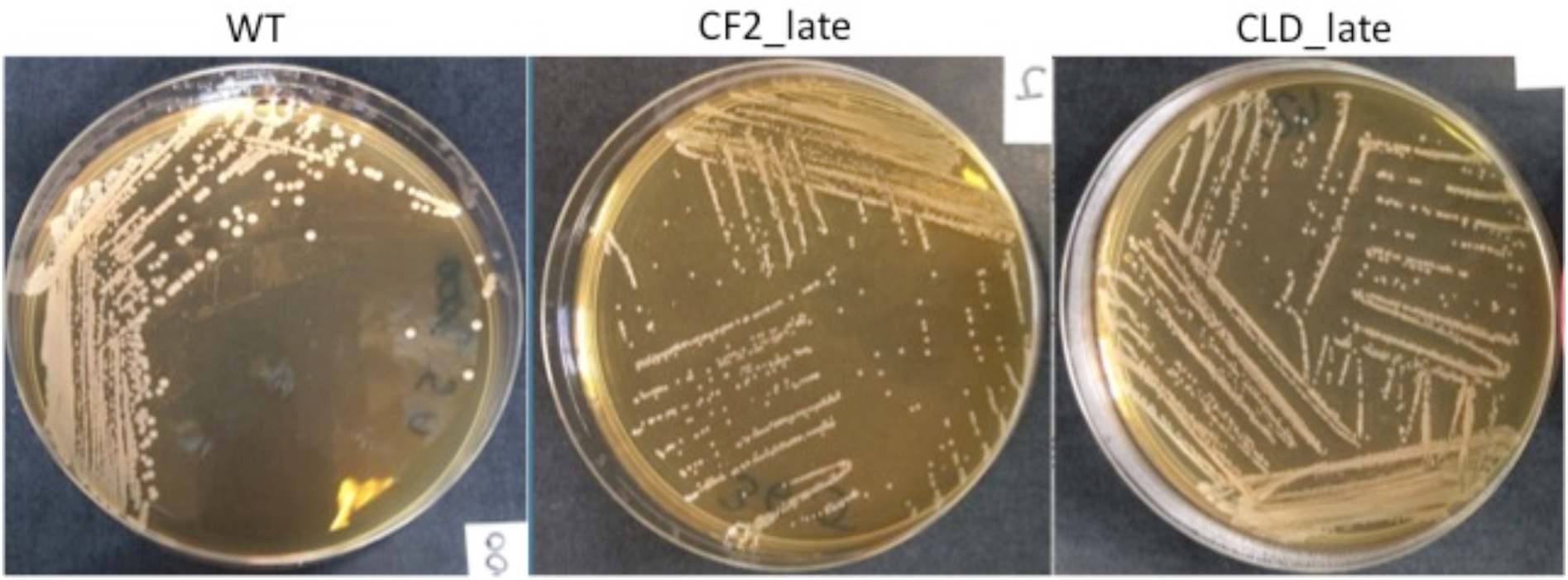
Colony morphology of USA300-LAC (WT), CF2_late and CLD_late isolates. Bacteria were grown on BHI agar incubated 12h at 37°C.

**Figure S5.**
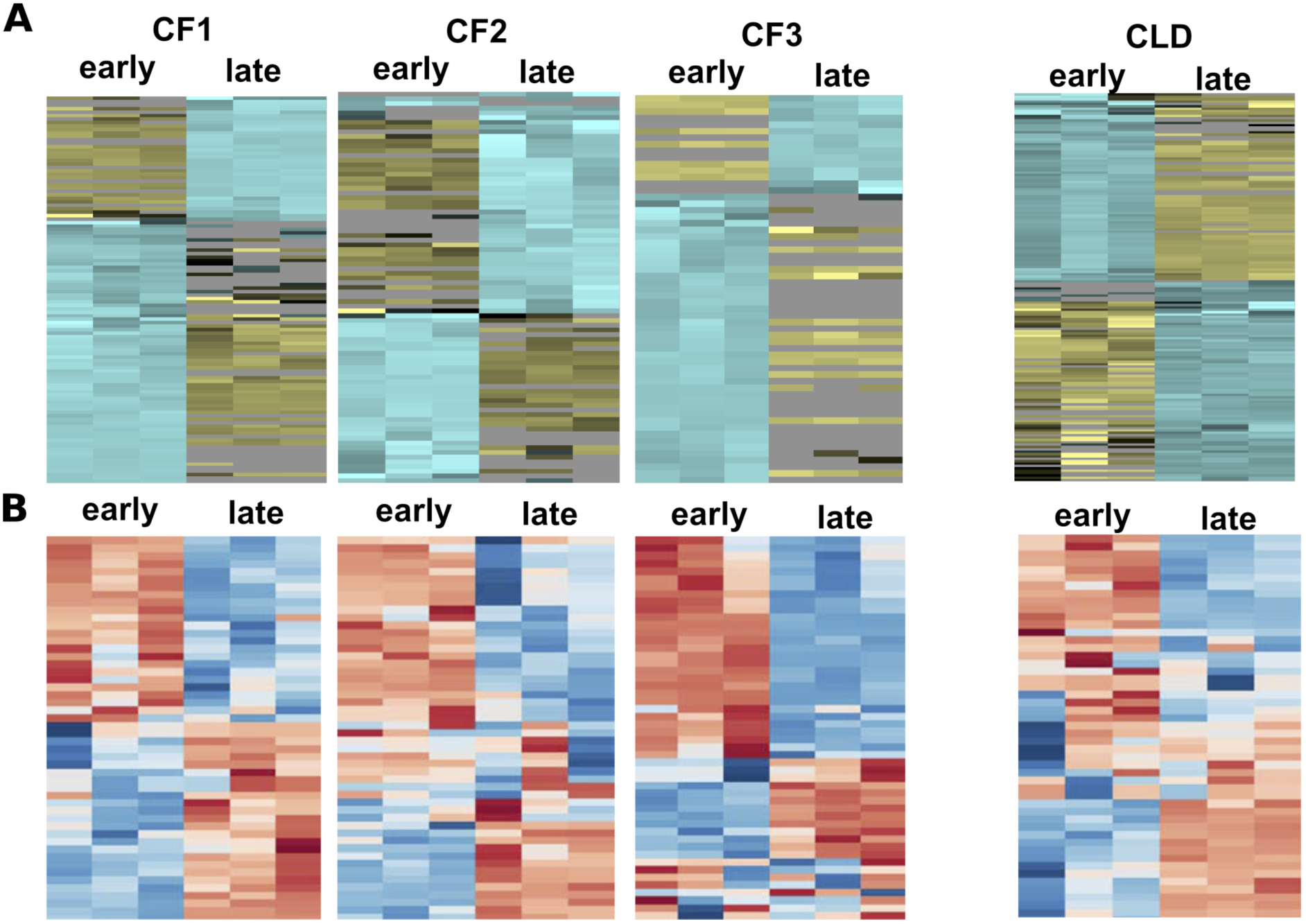
Heatmaps showing comparison of protein and metabolite profiles of early/late *S. aureus* isolates. The isolates were cultured to the stationary phase in medium mimicking the respiratory fluid of cystic fibrosis patients (CFSM) with the addition of thymidine. **A)** Heatmap visualization and hierarchical clustering analysis of the proteomic profiling in the late isolate compared to the early isolate of each patient. One experiment with three biological replicates was performed for each isolate. Rows: proteins; columns: samples; color key indicates protein relative concentration value (yellow: lowest; blue: highest). **B)** Heatmap visualization and hierarchical clustering analysis of the metabolite profiling in the late isolate compared to the early isolate of each patient. The top 50 most changing compounds are presented. Two independent experiments with three biological replicates were performed for each isolate. Rows: metabolites; columns: samples; color key indicates metabolite relative concentration value (blue: lowest; red: highest).

